# HIPPIE2: a method for fine-scale identification of physically interacting chromatin regions

**DOI:** 10.1101/634006

**Authors:** Pavel P. Kuksa, Alexandre Amlie-Wolf, Yih-Chii Hwang, Otto Valladares, Brian D. Gregory, Li-San Wang

## Abstract

Most regulatory chromatin interactions are mediated by various transcription factors (TFs) and involve physically-interacting elements such as enhancers, insulators, or promoters. To map these elements and interactions, we developed HIPPIE2 which analyzes raw reads from high-throughput chromosome conformation (Hi-C) experiments to identify fine-scale physically-interacting regions (PIRs). Unlike standard genome binning approaches (e.g., 10K-1Mbp bins), HIPPIE2 dynamically calls physical locations of PIRs with better precision and higher resolution based on the pattern of restriction events and relative locations of interacting sites inferred from the sequencing readout.

We applied HIPPIE2 to *in situ* Hi-C datasets across 6 human cell lines (GM12878, IMR90, K562, HMEC, HUVEC, NHEK) with matched ENCODE and Roadmap functional genomic data. HIPPIE2 detected 1,042,738 distinct PIRs across cell lines, with high resolution (average PIR length of 1,006bps) and high reproducibility (92.3% in GM12878 replicates). 32.8% of PIRs were shared among cell lines. PIRs are enriched for epigenetic marks (H3K27ac, H3K4me1) and open chromatin, suggesting active regulatory roles. HIPPIE2 identified 2.8M significant intrachromosomal PIR–PIR interactions, 27.2% of which were enriched for TF binding sites. 50,608 interactions were enhancer–promoter interactions and were enriched for 33 TFs (31 in enhancers/29 in promoters), several of which are known to mediate DNA looping/long-distance regulation. 29 TFs were enriched in >1 cell line and 4 were cell line-specific. These findings demonstrate that the dynamic approach used in HIPPIE2 (https://bitbucket.com/wanglab-upenn/HIPPIE2) characterizes PIR–PIR interactions with high resolution and reproducibility.

## Introduction

Enhancers are non-coding DNA elements that regulate gene expression by recruiting transcription factors which in turn mediate physical interactions with the promoters of their target genes to increase transcription of those genes. The genome-wide relationship between enhancers and their target genes depends on the three-dimensional DNA looping associated with enhancer–promoter interactions. To capture genome-wide chromatin interactions in Hi-C (Lieberman-Aiden et al., 2009), physically-interacting DNA regions and their binding proteins are cross-linked, followed by restriction enzyme cleavage and proximity ligation of the interacting DNA fragments to localize and capture pairs of interacting DNA fragments. These ligated DNA fragments are then sequenced to identify the chromatin interaction map genome-wide. Higher resolution in localizing interacting DNA fragments has been achieved by using a restriction enzyme with more frequent sites throughout the genome (e.g., MboI, a 4-cutter with a 4 base-pair motif, instead of a 6-cutter such as HindIII or NcoI) and by performing the DNA–DNA proximity ligation in intact nuclei to generate denser Hi-C contact matrices (Rao et al., 2014).

Previous methods for analyzing Hi-C data (Ay et al., 2014; Consortium et al., 2015; Durand et al., 2016; Forcato et al., 2017; Imakaev et al., 2012; Jin et al., 2013; Kaplan and Dekker, 2013; Lieberman-Aiden et al., 2009; Lun and Smyth, 2015; Ma et al., 2015; Norton et al., 2018; Rao et al., 2014; Yaffe and Tanay, 2011; Yang et al., 2018) have implemented a binning-based scheme for identifying interacting genomic regions, where reads are aggregated into equally-sized bins by genome coordinates and interacting regions are identified as pairs of bins with significant enrichments of reads using statistical models accounting for biases (e.g. negative correlation between linear genomic distance, number of reads and mappability) of the individual bins. While binning is effective at delineating large-scale chromatin structure, it does not capture specific physically-interacting DNA regions. The methods of Jin et al. and Hwang et al. (Hwang et al., 2013, 2014; Jin et al., 2013) have shown that it is possible to study interactions at the level of restriction fragments (i.e. the DNA region between two consecutive restriction sites) rather than bins using 6-cutter restriction. Restriction fragment-based binning might be problematic for more frequent cutters such as 4-cutter (e.g. MboI) (Rao et al., 2014), since restriction fragment length is much smaller on average and interacting DNA sites are more likely to span more than one restriction fragment.

To address these limitations, we propose HIPPIE2 (**Figure 1a,b**), a novel computational method that infers the locations of DNA physically-interacting regions (PIRs) by identifying regions enclosed by restriction events observed on both sides of DNA-protein Hi-C construct (**Figure 2**, Methods; **Supplementary Figure 6**). This strategy allows HIPPIE2 to identify individual interacting DNA elements with better specificity than binning. HIPPIE2 uses cell type-matched functional genomics data to characterize the interacting PIRs into functional categories including enhancers and promoters. This enables the high-resolution identification of cell type-specific enhancer–promoter interactions, and we show a corresponding enrichment in PIR–PIR interactions of transcription factor binding sites (TFBSs) for transcription factors known to be involved in enhancer–promoter interactions. HIPPIE2 is open source (https://bitbucket.com/wanglab-upenn/HIPPIE2) and freely available as a full pipeline to automate analysis from raw Hi-C reads to identification of PIRs, significant PIR–PIR interactions, functional genomics annotations and TF analysis.

**Figure 1.**
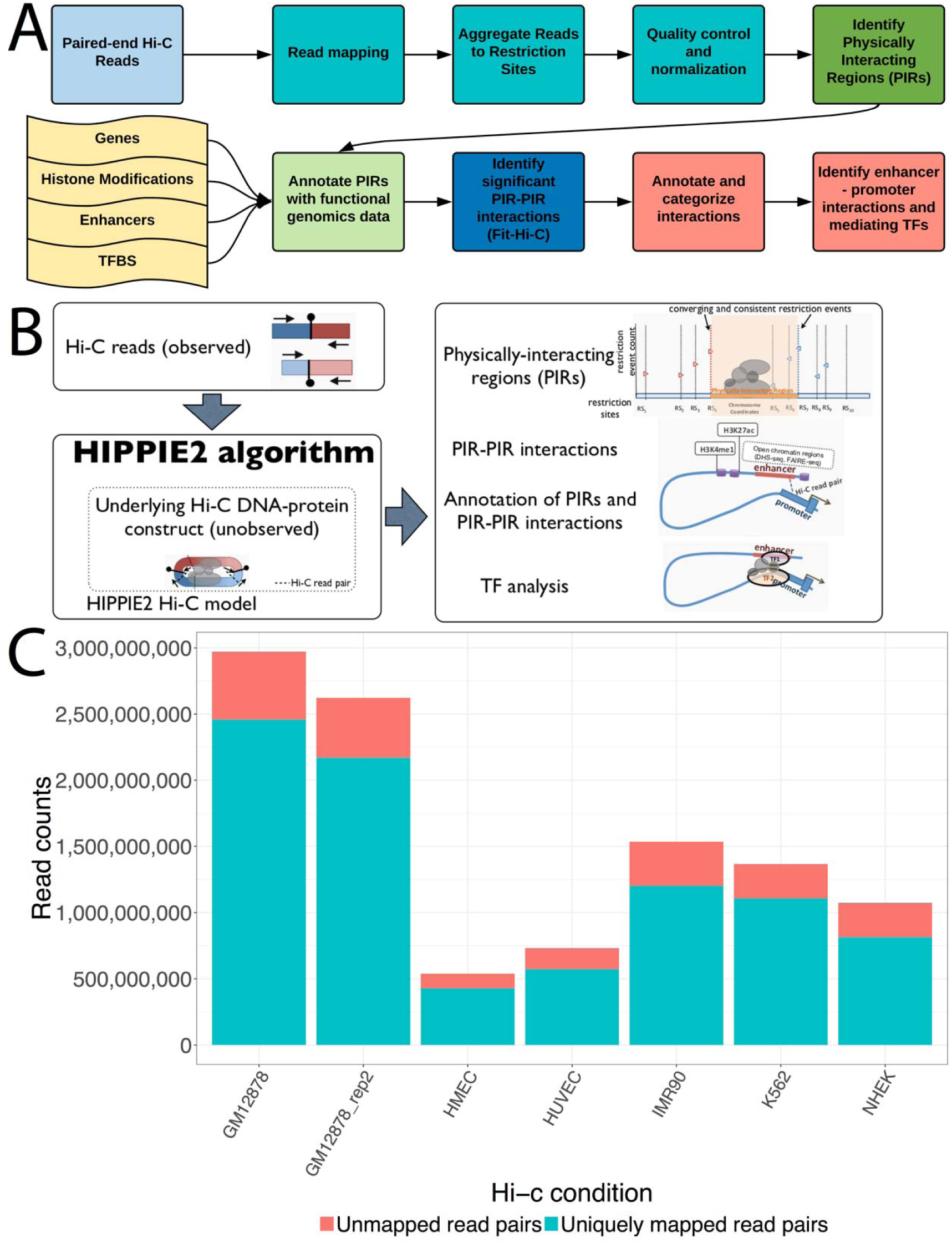
Description of HIPPIE2 pipeline and mapping statistics. A) Detailed processing pipeline of HIPPIE2. B) Overview of HIPPIE2 algorithm. C) Mapping statistics across cell lines.

**Figure 2.**
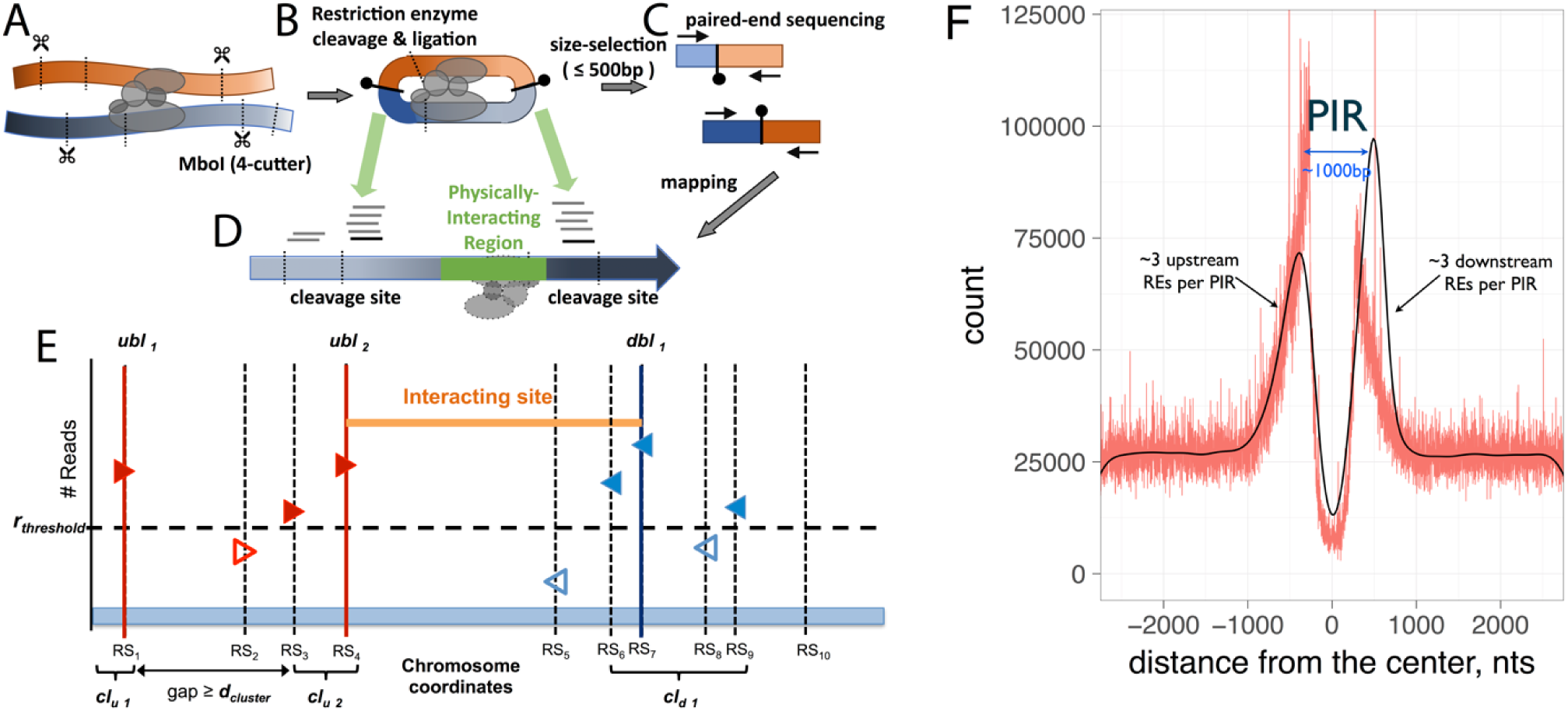
Hi-C model and identification of DNA physically-interacting region. A) Interacting DNA regions are cut by the MboI restriction enzyme. B) Cut fragments are ligated together and size-selected. C) Paired end-sequencing is performed on the ligated fragments. D) Read pileups around the cleavage sites inform the identification of the physically-interacting region. E) Genomic view of read pileups, restriction sites, and interacting regions (PIRs) locations. Upstream (ubl) and downstream (dbl) boundary locations for PIRs correspond to most consistently cut (as evidenced by the number of reads) restriction/ligation sites. F) Distribution of restriction events (REs) around physically-interacting DNA regions (PIRs) identified by HIPPIE2. Shown is the distribution of the restriction events for PIRs in GM12878 cell line.

## Results

### HIPPIE2 identifies fine-scale physically-interacting DNA regions (PIRs)

The HIPPIE2 method presented in this manuscript further develops our HIPPIE method (Hwang et al., 2014): HIPPIE2 applies a newer read mapping protocol to resolve chimeric reads, interaction calling algorithm, and introduces novel algorithms to dynamically identify fine-scale interacting regions (**Figures 1, 2**) instead of binning reads into full restriction fragments used in HIPPIE (Hwang et al., 2014). To illustrate our method, we applied HIPPIE2 to analyze high read depth Hi-C sequencing datasets (Rao et al., 2014) using the 4bp-cutter MboI across six human cell lines that had matching functional genomics data from ENCODE or Roadmap (Bernstein et al., 2012; Consortium et al., 2015) including K562, HMEC, HUVEC, IMR90, NHEK, and GM12878, with two replicates for GM12878 (**Figure 1a-b**). For each cell line, we mapped the raw Hi-C read-pairs using STAR (Dobin et al., 2013) (**Methods**), uniquely mapping between 73.6% and 85.4% of Hi-C reads across the 51 separate libraries for these cell lines (**Figure 1c, Supplementary Figure 1**). Following (Rao et al., 2014), we normalized the read counts using matrix normalization by Knight and Ruiz (Knight and Ruiz, 2012). Using normalized counts, HIPPIE2 identifies physically-interacting regions (PIRs) as the DNA regions flanked on both sides by restriction sites (RSs) that were observed to be consistently cleaved/ligated in a given Hi-C sequencing library, using information from the Hi-C sequencing read-out including the read mapping coordinates, distances from reads to their nearest restriction sites, DNA ligation constraints and strand orientations (+/-) of the mapped read-pairs, and relative locations of DNA interaction sites with respect to mapped reads (Methods, **Figure 2**). This dynamic PIR-based approach enables finer-scale identification of specific interacting DNA regions compared to the binning approach (**Table 1**).

**Table 1:** Comparison of HIPPIE2 with other Hi-C methodologies

| Method | Input | Resolution | Output | Downstream Analyses |
| --- | --- | --- | --- | --- |
| HIPPIE2 | Raw reads | Dynamic, restriction event-based (1kb average) | High-resolution restriction site-based physically-interacting regions (PIRs), PIR-PIR interactions | Genic and cell type-specific epigenetic annotation of interactions, identification of mediating transcription factors, identification of enhancer-promoter interactions |
| HIPPIE (Hwang et al., 2014) | Raw reads | Restriction fragment-based (4kb average) | Full restriction fragment-based physically-interacting regions | Annotated interactions, enhancer-promoter interactions |
| HiCCUPS (Durand et al., 2016; Rao et al., 2014) | Raw reads | Fixed bins | loops (bin-bin interactions) | NA |

In total, HIPPIE2 called between 1,584,000 and 1,886,000 PIRs from chromosomes 1-22 and X across cell lines (**Figure 3a**). These PIRs had an average length of 1,006 base pairs consistent across cell lines (**Supplementary Figure 2a**), which corresponds to 2.4 average restriction fragment length. Across libraries, these identified PIRs covered 53.2%-59.3% of the genome (**Supplementary Figure 2b**). HIPPIE2 annotated these PIRs with gene annotations (Pruitt et al., 2014) including promoters, exons, and introns, which found that a majority of PIRs in all cell types were intergenic and the next largest class of overlaps were in mRNA introns, supporting the regulatory roles of these PIRs (**Figure 3b**). Comparing with DNase-seq-based regions of open chromatin in the matching cell types from Roadmap/ENCODE, we found that 73.84-79.04% of open chromatin regions were covered by PIRs across cell types, with an average of 69.97% of the open chromatin peaks covered by PIRs (**Figure 3c**). PIR identification by HIPPIE2 is highly robust, with 92.3% PIRs (1,649,417) found in the two GM12878 replicates.

**Figure 3.**
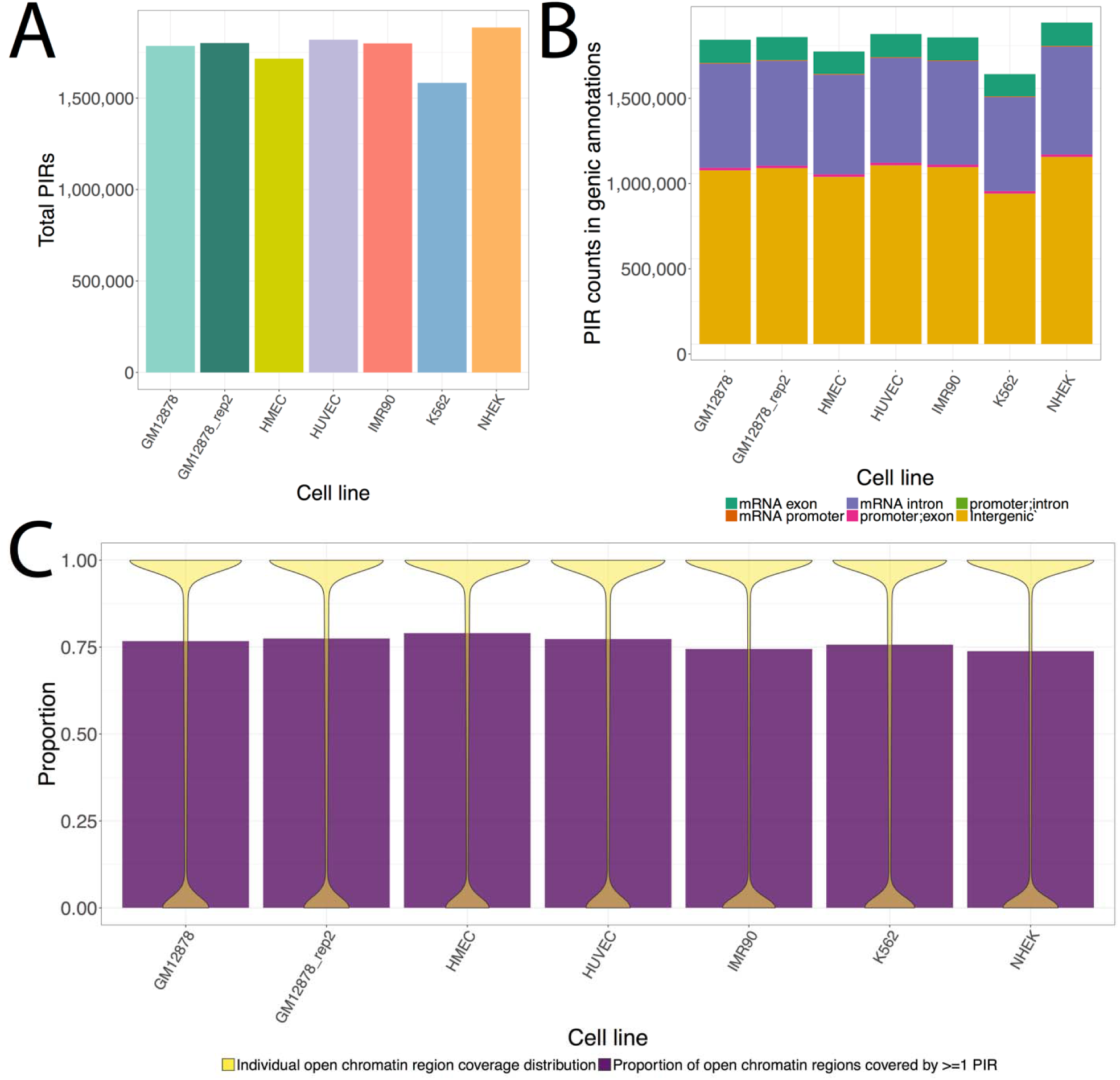
PIR characteristics. A) Total PIRs identified across cell lines. B) Localization patterns of PIRs in various genomic annotations. C) Overlap patterns of PIRs with cell type-matched open chromatin annotations. Yellow distributions display the proportions of individual open chromatin peaks by PIRs and purple bars are proportion of all open chromatin peaks with any PIR coverage.

### HIPPIE2 detects fine-scale chromatin interactions

To identify which PIR–PIR pairs are significantly interacting, HIPPIE2 applies Fit-Hi-C algorithm (Ay et al., 2014) using the normalized read counts and linear genomic distance between pairs of potentially interacting PIRs (Methods). Across cell lines, HIPPIE2 identified between 42,500 and 1,194,010 intra-chromosomal significant PIR–PIR interactions (>5 kb apart, adjusted P value ≤ 0.05, **Figure 4a**). To investigate robustness of interaction calling, we compared the two GM12878 replicates. Consistent with the lower sequencing depth of the replicate library (2.5 billion vs 3 billion reads), we identified fewer significant interactions and PIRs involved in significant interactions in the replicate library (**Figure 4a**), but found a significant overlap between PIRs and PIR–PIR interactions (**Figure 4b-c**), with majority (66.2%; 274,445/414,343) of PIRs and 31.1% (196,343/631,610) of PIR–PIR interactions in the replicate found in the primary library. This level of replication is consistent with prior studies of Hi-C replication (Forcato et al., 2017), which found similar levels of replicated interactions across Hi-C datasets.

**Figure 4.**
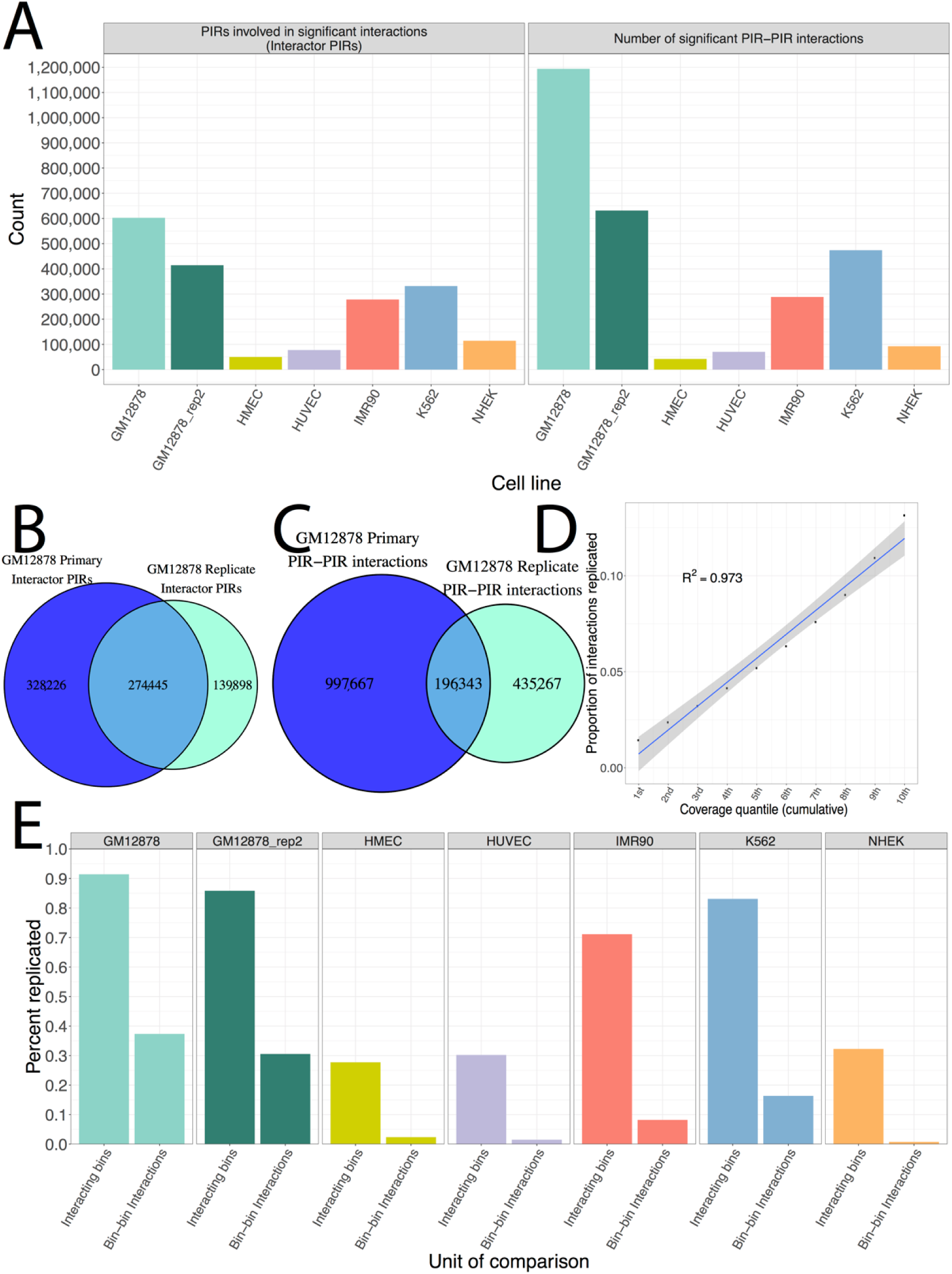
Characteristics of significant PIR–PIR interaction identification and replication. A) Counts of PIRs involved in significant interactions (left) and number of significant PIR–PIR interactions (right) across cell lines. B) Number of PIRs involved in significant interactions replicated between the primary and secondary GM12878 libraries. C) Number of PIR–PIR interactions replicated between the primary and secondary GM12878 libraries. D) Plot of replication rate against PIR read coverage quantiles. Correlation is the Pearson correlation. E) Replication of Rao data by HIPPIE2. Interacting bins refer to 10kb Rao bins involved in significant interactions and bin–bin interactions are significant interactions.

To interrogate the relationship between sequencing depth and interaction replication, we binned the interactions from the GM12878 primary library into deciles by read coverage (analogous to down-sampling) and compared their replication rates. We found a striking positive correlation between read coverage and replication rate (R^2^ = 0.9398, **Figure 4d**), suggesting that reproducibility between replicates may be increased with a higher sequencing depth.

### Comparison with uniform binning-based approach

HIPPIE2 provides a more accurate approach for identifying fine-scale interacting sites by design: previous methods that use a binning-based approach maximize their statistical power to detect interactions, with a tradeoff of accuracy for identifying the interacting site (**Table 1**). Due to the fundamentally different natures of the binning-based algorithms compared to HIPPIE2 and the lack of a ‘ground truth’ dataset of expected Hi-C interactions, it is challenging to directly compare the HIPPIE2 interactions with these previous methods. However, to explore the differences between the binning approaches and HIPPIE2 PIR-based approach, we compare the HIPPIE2 results with the results from (Rao et al., 2014) obtained using HiCCUPS method (we note that a detailed comparison among Hi-C methods has been reported in the recent study by Forcato et al (Forcato et al., 2017)).

We compared our HIPPIE2-identified PIRs with the HiCCUPS-identified loop anchors and interactions (bin size=10K) (Rao et al., 2014). Across cell lines, the set of PIRs identified by HIPPIE2 is consistent with and is complementary to the previously identified set of interacting genomic regions: we found that HIPPIE2 PIRs covered an average of 60.2% of HiCCUPS-identified loop anchors across cell lines, with the highest proportion (91.4%) in the primary, most deeply sequenced GM12878 library (Methods, **Figure 4e**). The HMEC, HUVEC, and NHEK cell lines were the only ones with a proportion less than 50%, corresponding to their shallower sequencing depth (Pearson R^2^ = 0.862 between sequencing depth and replication proportion).

Next considering HIPPIE2 PIR–PIR interactions and HiCCUPS loops, we found that 37.4% of HiCCUPS loops in GM12878 primary library, with an average of 16.6% of HiCCUPS loops across cell lines, were supported by significant HIPPIE2 PIR–PIR interactions across cell lines (**Figure 4e**). When we matched the bin size (10k) used in HiCCUPS analysis by expanding HIPPIE2 PIR–PIR interactions so that each PIR covered at least 10kb, we found that the majority (55.8%) of HiCCUPS loops replicated in the primary GM12878 library (highest sequencing depth) and the average proportion of replicated HiCCUPS loops across cell lines increased to 28.88% from 16.6%, with an average of 68% of HiCCUPS loop anchors replicated (**Supplementary Figure 3c**). Interestingly, each PIR overlapping a HiCCUPS-identified loop anchor was involved in an average of between 5.62 and 21.52 significant PIR–PIR interactions across cell lines compared to a single interaction (loop) reported by HiCCUPS, corresponding to an average fold enrichment of 10.07 more interactions identified by HIPPIE2 than by the HiCCUPS binning/loop detection approach. Overall, HIPPIE2 identified about two orders of magnitude more interactions than the bin-based approach across all cell lines (2,794,123 vs 27,827). This illustrates that HIPPIE2 identifies more, finer-scale regulatory interactions than the bin-based approach to maximize power to detect large-scale genomic architecture rather than a multiplicity of fine-scale regulatory interactions. For example, in the 1 megabase locus on chr14 investigated in Rao et al (Rao et al., 2014) (chr14:94,000,000-95,000,000), there were 1,082 HIPPIE2 significant PIR–PIR interactions within 4 bin-bin interactions that were called in the original study (**Supplementary Figure 4**).

### HIPPIE2 PIR–PIR interactions are enriched in regulatory genomic features

To evaluate how PIRs co-locate with binding of factors known to be involved in genome architecture and transcriptional mechanisms, we overlapped HIPPIE2-identified PIRs from the primary GM12878 library, the most deeply sequenced library, with ENCODE ChIP-seq binding sites for CTCF, PolII, and P300 (**Table 2**) from the same cell type (Bernstein et al., 2012). We found that HIPPIE2 PIRs overlapped 92% (41,134 out of 44,597) CTCF sites, consistent with studies that suggests CTCF has a role in mediating chromatin interactions (Phillips and Corces, 2009). Similarly, for PolII and P300, associated with transcriptional and enhancer activity (Shlyueva et al., 2014), we found high overlaps at 96.5% (9,678 out of 10,026) and 96.3% (16,509 out of 17,150 sites), respectively. We randomly sampled 1,000 sets of background genomic regions matched to the distribution of PIR lengths and calculated percent enrichment of the GM12878 PIRs relative to these background sites (Methods, **Table 2**). We found that the GM12878 PIRs had increases of 20-23% base pairs of overlap over background and overlapped 18-19% more ChIP-seq sites, suggesting that the GM12878 PIRs are involved in genomic architecture and regulatory function.

**Table 2:**
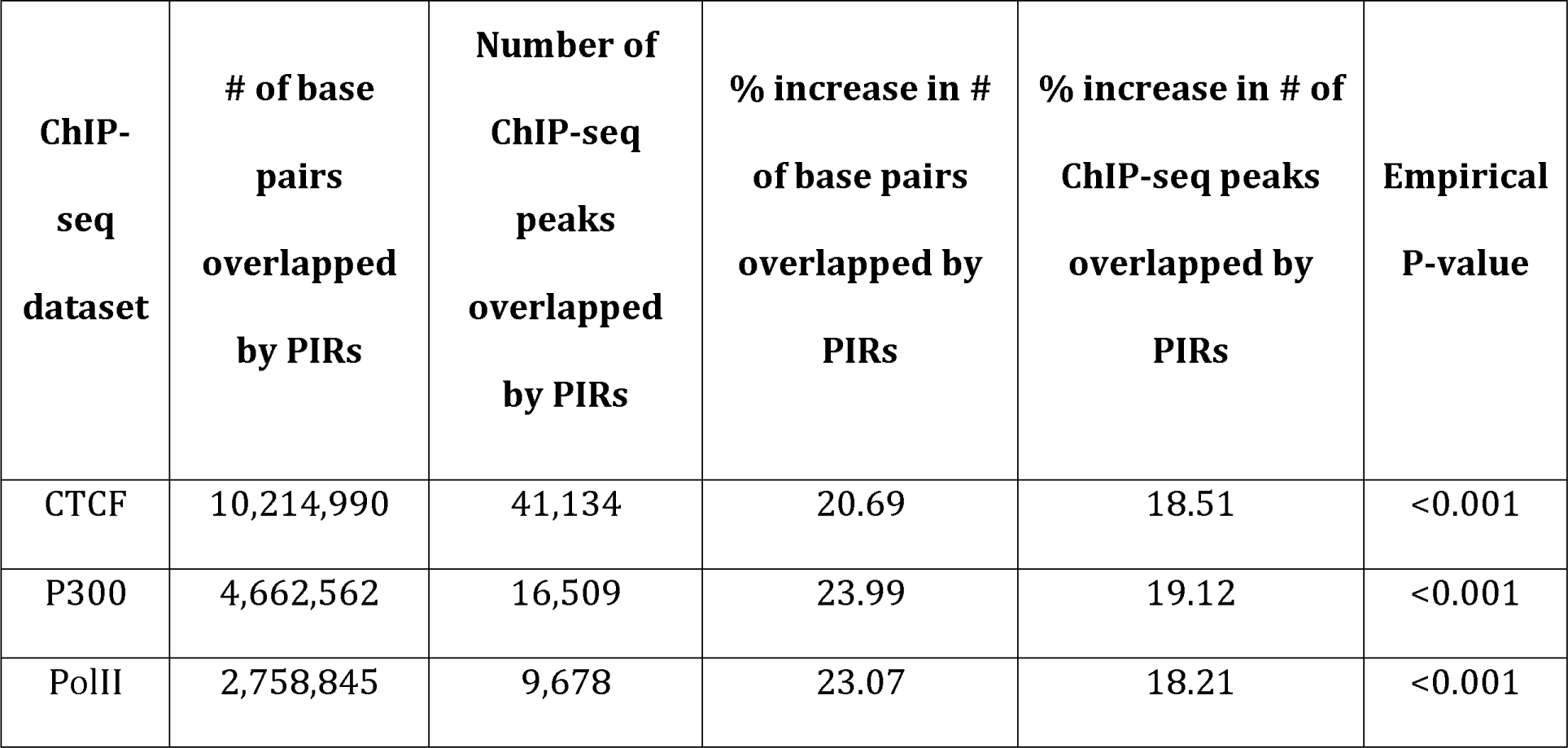
Enrichment of all GM12878 PIRs in transcriptional and architectural protein binding sites relative to randomly sampled background genomic regions

To characterize the function of PIRs involved in significant interactions, HIPPIE2 automatically annotates PIRs with DNase-based open chromatin regions, enhancers defined by combinatorial epigenomic status using ChromHMM (Ernst and Kellis, 2012), the enhancer-associated histone modifications H3K4me1 and H3K27ac (Calo and Wysocka, 2013; Thurman et al., 2012), the inactive or poised enhancer histone modification H3K27me3 (Zhu et al., 2013), and the promoter-associated histone modification H3K4me3 (Bernstein et al., 2006), all in the matching cell types from ENCODE or Roadmap (Bernstein et al., 2012; Consortium et al., 2015) (**Figure 5a**,**b**). Between 18.35% and 42.26% of PIRs involved in significant PIR–PIR interactions overlapped open chromatin sites across cell lines, with the lowest proportions in the shallowest sequencing libraries, while the other annotations encompassed between 7.4% (H3K4me3) and 17.2% (Roadmap ChromHMM enhancers) of PIRs on average across cell lines. By comparing against samples of length-matched background intervals (Methods), we found that the PIRs involved in significant interactions were enriched for overlaps with all of the active epigenomic marks in every cell line except for NHEK, where PIRs were depleted of overlaps with all annotations except H3K4me3 (**Figure 5b**). The repressive mark H3K27me3 had the smallest average enrichment (40.75%) across cell lines, suggesting that significant PIR interactions are associated with active regulatory elements (Hwang et al., 2013; Rao et al., 2014), which showed much stronger enrichments.

**Figure 5.**
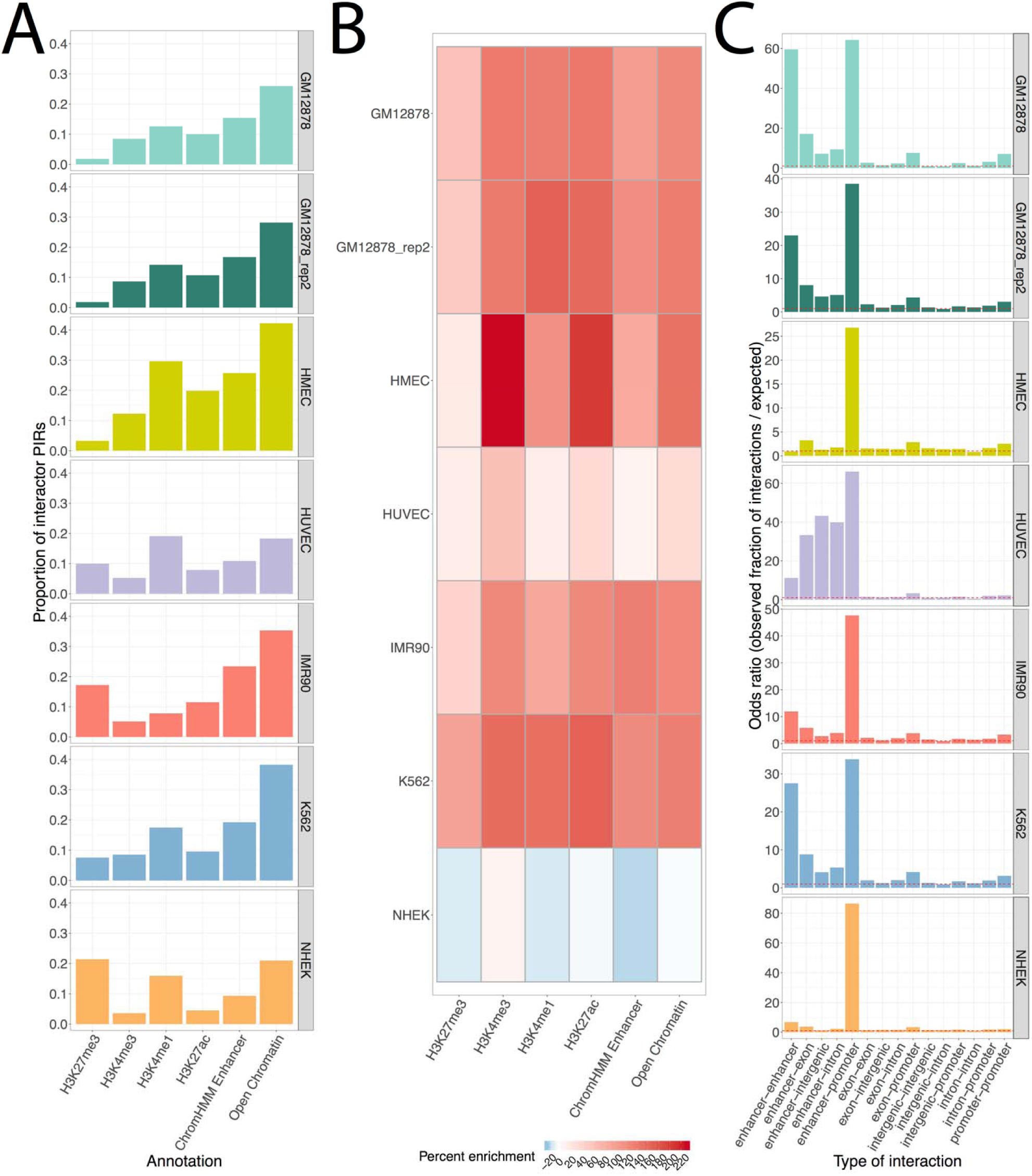
Regulatory annotation of PIR–PIR interactions. A) Proportion of PIRs involved in significant interactions (interactor PIRs) overlapping cell type-matched functional annotations. B) Enrichment of interactor PIRs relative to background expectation calculated by sampling. C) Ratio of number of observed annotation-annotation PIR–PIR interactions relative to background expectations.

### HIPPIE2 PIR–PIR interactions are enriched for enhancer–promoter mechanisms

HIPPIE2 identifies specific enhancer–promoter interactions by classifying PIRs as enhancers if they overlap 1) an open chromatin region and a shared H3K4me1/H3K27ac peak and/or 2) a ChromHMM epigenomic enhancer (Methods). For enhancer elements, HIPPIE2 further requires that the putative enhancer PIR display at least one significant interaction with a promoter-overlapping PIR and does not overlap any H3K4me3 (active promoter mark) or H3K27me3 (repressive mark) peaks in the matching tissue. We found that the percentage of regulatory interactions (enhancer–promoter, enhancer–enhancer, or promoter–promoter pairs) accounted for an average of 2.36% (ranging from 0.36% - 3.78%) of significant interactions across cell lines, a significant enrichment compared to the background expectation of an average of 0.076% (ranging from 0.0049% - 0.2%) of interactions. For enhancer–promoter interactions specifically, we detected an average of 51.95x enrichment over the background expectation (ranging from 26.79x - 86.47x), and these were the most enriched interactions in all cell types, suggesting that the interactions identified by HIPPIE2 are indeed reflective of transcriptional regulatory processes (Methods, **Figure 5c**).

### HIPPIE2 recovers a repertoire of known regulatory TFs mediating chromatin interactions

In order to elucidate the mechanisms underlying the observed enhancer–promoter interactions, HIPPIE2 annotates interacting PIRs with transcription factor binding sites (TFBS) from the FactorBook database which contains TFBSs for 133 DNA-binding proteins identified by ChIP-seq experiments (Wang et al., 2013). Combined with our HIPPIE2-identified fine-scale PIR annotations, this approach enables HIPPIE2 to identify the transcription factors mediating enhancer–promoter interactions with high resolution. We found that an average of 14.11% of PIRs involved in all significant interactions had overlaps with known TFBS across cell lines, while an average of 39.2% of HIPPIE2-identified enhancer–promoter interactions across cell lines had an evidence of known TF binding (**Supplementary Figure 5a**).

To determine whether enhancer–promoter interactions were enriched in transcription factor binding sites, we quantified the observed/expected ratio for binding motif enrichment and used a binomial model to identify significant enrichments of transcription factors involved in enhancer–promoter interactions (Methods). We found significant enrichments for 31 TFs in enhancers and 29 in promoters for a total of 33 unique transcription factors across all cell lines except for HUVEC and NHEK (**Supplementary Figure 5b, Supplementary Table 3**). To test whether these putative HIPPIE2-identifed TF–TF interactions correspond to known protein-protein interactions (PPI), we compared HIPPIE2 TF-TF interactions to the BioGRID database (Chatr-Aryamontri et al., 2015; Tyers et al., 2006). To do this, for each cell line, we identified all the TFs involved in significant enhancer–promoter interactions, quantified all their interactions in BioGRID, and determined the proportion of BioGRID interactions involving these TFs that were recapitulated by HIPPIE2. We found that HIPPIE2-identified TF–TF interactions in GM12878 recapitulated most (78%) of known physical TF–TF interactions reported in BioGRID, with an average of 57.5% of known physical interactions between TFs in BioGRID (31%-78%) recovered across cell lines. These proportions were strongly correlated with the sequencing depth of each cell line (Pearson R^2^ = 0.88), suggesting that increased read depth may recover more BioGRID interactions.

We then stratified these TFs by how many different cell lines they were enriched in to identify regulatory mechanisms common across cellular contexts (**Supplementary Figure 5c**). This identified several TFs enriched in several cell lines that were consistent with known enhancer and chromatin architecture biology, including SP1, AP1, MYC, CEBPB, YY1, and CTCF. SP1 has been shown to function as a link of both side of DNA, and is able to form a tetrameric structure and assemble multiple tetramers that facilitate a DNA looping structure (Mastrangelo et al., 1991). AP1 is a transcription factor involved in cellular proliferation, transformation, and apoptosis that forms heterodimers with the Jun oncogene (Shaulian and Karin, 2002). MYC is an oncogene involved in several different cancer types and exerts widespread transcriptional regulatory effects (Dang, 2012). CEBPB is another major enhancer-binding protein family which can aid the transition of enhancer elements from closed chromatin to a primed or poised state and is involved in immune and inflammatory responses (Heinz et al., 2015). CTCF is a major architectural protein with a role in defining megabase-scale topologically-associated domains as well as regulating smaller-scale enhancer–promoter interactions such as those observed here (Ong and Corces, 2014; Phillips and Corces, 2009). YY1 is another major architectural protein that cooperates with CTCF to mediate looping interactions involved in developmental processes and enhancer–promoter interactions (Beagan et al., 2017; Weintraub et al., 2017).

## Discussion

In this paper we introduce a novel method for Hi-C data analysis, HIPPIE2, which dynamically discovers fine-scale physically-interacting regions (PIRs) of the genome with increased resolution compared to previous methods, detects fine-scale chromatin interactions, and provides functional and mechanistic characterization of these interactions. HIPPIE2 uses the pattern of restriction events as evidenced by sequencing read pileups relative to restriction sites to fine-map interacting DNA regions. Our results suggest that HIPPIE2 detects more specific, finer-scale interactions at the gene-regulatory level of chromatin architecture (average PIR length of 1,006bps), offering a complementary approach to the binning-based procedures (Ay et al., 2014; Durand et al., 2016; Forcato et al., 2017; Heinz et al., 2010; Mifsud et al., 2015; Rao et al., 2014). Our method also complements restriction fragment-based methods (Hwang et al., 2014; Jin et al., 2013) as an alternative for analyzing data from more frequent cutters with much smaller fragment length and interaction regions spanning more than one fragment.

With our approach designed to work at the inherent resolution of the data (as determined by restriction enzyme cutting frequency and restriction efficiency), our method will prove useful in the analysis of chromosome conformation capture experiments with further increased sequencing depth or improved restriction protocols. Another natural application in which our approach will prove useful is the analysis of the data generated by the assays targeting particular types of interactions, such as Capture-C and Capture Hi-C (Hughes et al., 2014; Mifsud et al., 2015) that capture promoter-centric interactions.

Furthermore, the fine-scale resolution of our method for detecting interacting regions enables analysis, identification and interpretation of specific proteins/TF complexes mediating these interactions. This TF analysis can be improved by *de novo* motif discovery in PIR sequences, incorporation of protein–protein interaction networks to identify protein/TF complexes, and protein domain compatibility information. Using the identified interacting sequences and mediating TFs can help build predictive models for regulatory interactions such as (Schreiber et al., 2018; Whalen et al., 2016). Another direction in which our fine mapping HIPPIE2 method will prove useful is in comparison and analysis of changes in fine-scale regulatory networks during development or between different conditions. HIPPIE2 is freely available as an open source pipeline (https://bitbucket.org/wanglab-upenn/HIPPIE2). HIPPIE2 generated interaction data is also available in UCSC Genome Browser hub (https://genome.ucsc.edu/s/alexamlie/HIPPIE2%20vs%20Rao%20all%20cell%20lines%20darker%20interaction%20lines).

## Methods

### Hi-C data acquisition and genome mapping

For our analysis, we used the Hi-C datasets for GM12878 (primary and secondary replicates), HMEC, HUVEC, IMR90, K562, and NHEK cell lines from (Rao et al., 2014), acquired from the GEO database (accession number GSE63525). For each condition, we acquired FASTQ files from the SRA files available on GEO corresponding to sequencing libraries within each condition. Each library was mapped separately and then combined for downstream analyses. HIPPIE2 first aligns the paired-end reads to the human genome (hg19 assembly) using the STAR aligner (Dobin et al., 2013) allowing only unique mapping (full parameters available in HIPPIE2 open source repository, starMappingToBam.sh script). Each of the single-end reads from a read-pair was first mapped separately and then re-associated with the corresponding second read in a read-pair. To improve mapping, both contiguously mapped and chimeric reads are identified and paired. Both halves of a chimeric read were required to map uniquely and have a minimum mapped length of 22 nt. For those paired-end reads with a chimeric read involved, we required that the pairing partner of the chimeric read (a single-end read) mapped in the proximity of one of the two split halves spanned by the chimeric read.

### Hi-C read normalization

To remove potential random ligation events, including un-cut, self-ligated, or re-ligated read-pairs, we filtered out the read-pairs that are less than 5,000 bps apart from each other as suggested in (Jin et al., 2013; Lajoie et al., 2015). In addition, to correct for all possible Hi-C experimental biases including length of the crosslinked DNA fragments, restriction site accessibility, or ligation rate of the restriction enzyme digested fragments, we normalized the read counts using the matrix normalization method by Knight and Ruiz (Knight and Ruiz, 2012) as used in (Rao et al., 2014). Additionally, to avoid any biases on detecting the region that cannot be mapped as a unique genomic locus, we also removed from the analysis restriction sites (RSs) that have mappability less than 0.8. We found that 96% of the RSs have mappability higher than 0.8, i.e. most of RSs had high mappability given a relatively long read length (101 nts).

### Identification of physically-interacting regions

To identify physically-interacting DNA regions (PIRs), we utilized the idea that each single-end Hi-C read is always located in the proximity of a restriction site (RS) that serves as both the restriction enzyme cleavage and ligation site in the Hi-C protocol. The RSs correspond to sites in the genomic DNA containing sequence that can be recognized by the restriction enzyme, e.g., “GATC” for restriction enzyme MboI. After HIPPIE2 maps reads, it first determines corresponding RSs (cleavage/ligation sites), and infers the relative position (upstream or downstream from the RSs) for the DNA-interacting region (PIR).

The cleavage/ligation sites are identifiable from the mapping information of Hi-C paired-end reads because (1) a proper DNA ligation forms a phosphodiester bond between the 5’ phosphate of the donor DNA and the 3’ hydroxyl of the acceptor DNA, and (2) the strand orientation pattern reported by Illumina sequencer is restricting the combinations of upstream or downstream cleavage/ligation site of each read-pairs. The workflow of identifying all physically-interacting regions (PIRs) and PIR–PIR interactions along the genome includes three major phases: (I) find ligation junctions for read-pairs (II) identify physically-interacting regions, and (III) find PIR–PIR interactions. Each phase is described below.

#### i. Identifying ligation junctions for read-pairs

Each mapped read has two candidate (nearest upstream or downstream) restriction sites (RSs) to be assigned as the restriction enzyme cut-and-ligation site. To determine the cut-and-ligation site, we first determine which type of interaction has happened based on the mapped strand orientation (**Supplementary Figure 7, Supplementary Figure 6a, Supplementary Figure 8**). We enumerated all the possible ligations types: head/tail, tail/head, head/head, or tail/tail ligations; where head is the end with smaller genome coordinate and tail is the end that is on a larger coordinate of the chromosome.

Because (1) the ligation of two DNA fragments is formed by a phosphodiester bond between a 3’ hydroxyl and a 5’ phosphate, and (2) Illumina paired-end sequencing reads are generated from opposite strands from the sequenced DNA fragments, we can narrow down four possible ligation types for each paired-end reads to two scenarios using its strand orientation. For the read strand combinations of +/- or -/+ (different strand), the two possible ligation types are either head/tail or tail/head ligations (**Supplementary Figure 7 left**). Similarly, for the read strand combinations of +/+ or -/- (same strand), the two possible ligation types are either head/head or tail/tail ligations (**Supplementary Figure 7 right**). Next, because of the size-selection step in the Hi-C protocol, cut-and-ligation events are expected to generate read pairs within 500bp of the restriction enzyme (MboI) cutting sites due to the size selection, to resolve the two possible cases of head/tail or tail/head for +/- and -/+, we calculated the two possible sums of the two distances to the nearest cutter sites, and ruled out the ligation event that made the sum larger than 300 base pair, which would be result from ligation of nonspecific cleavage product (Yaffe and Tanay, 2011)in the Hi-C experiment (**Supplementary Figure 8**). As shown in **Supplementary Table 2**, the observed fractions of strand orientation combinations for sequenced Hi-C read pairs are close to uniform as expected from the stochastic nature of the proximity ligation reaction.

#### ii. Identifying physically-interacting regions (PIRs)

With the identified RSs that form DNA-DNA ligation junctions, we further identify physically-interacting regions (Algorithm in **Supplementary Figure 6b**). First, we note the sum of upstream and downstream read counts (single-end reads from read-pairs) for each RS identified in the previous step. To group restriction events corresponding to the same interaction (interacting region), we clustered RSs separately for upstream and downstream read counts by thresholds of the maximum gap (*d*_*cluster*_) and the minimum read (*r*_*threshold*_). The maximum gap is defined as the third quantile of the restriction fragment distance distribution, and the minimum read requirement is defined as the median of the normalized read distribution for each chromosome. Within each corresponding cluster, we identify the RSs with the maximum read count (i.e. most consistently cut site) as the candidate flanking ends for PIRs. Finally, we matched the nearest upstream and downstream candidate flanking ends with a max-gap algorithm (in this study, the max-gap is 4000 bp), and report the PIRs as regions that are enclosed by the upstream and downstream RSs with the maximum read count in the upstream and downstream restriction clusters.

#### iii. Finding all PIR–PIR interactions

We find the interactions between PIRs by tracing the Hi-C read-pairs that participated in the identifications of PIRs (Algorithm in **Supplementary Figure 6c**). For each PIR identified in the previous step, IDs of single-end reads in the left and right RS clusters are used to identify PIRs containing mate reads (i.e. other single-end reads from read-pairs) as interacting partners. All such PIR–PIR interactions are then reported along with the read counts.

### Identifying significant PIR–PIR interactions genome-wide

To identify significant intra-chromosomal PIR–PIR interactions, we applied the Fit-Hi-C method (Ay et al., 2014) in R v3.2.3. For each of the autosomal chromosomes (1–22) and chromosome X, we split all observed PIR–PIR interactions into 2,000 distance groups according to the linear distance (in nucleotides) between interacting PIRs. We filtered out the PIR pairs that are less than 5,000 nucleotides apart. For each distance group, we calculated the average distance and the average normalized read counts of the interacting PIRs. With the 2,000 aggregated data points, we fit the normalized read counts by the function of distance using *smooth.spline* function in *R*. After the first spline fitting, we removed the outliers as described in (Ay et al., 2014) and fit the second spline function. We then reported PIR–PIR interactions that are significant after Benjamini–Hochberg correction (adjusted *P* values <= 0.05).

### Overlap with DNA loops from Rao et al

We compared PIR–PIR interactions in our study with the set of DNA loops identified in (Rao et al., 2014) using HiCCUPS. We downloaded the set of loops and Hi-C loci (DNA regions that are participating in significant DNA loops) and filtered to include only those with the highest 10 kb resolution from GEO database under accession number GSE63525. We then overlapped these loop anchors and interactions with HIPPIE2-identified PIRs and PIR-PIR interactions using a custom script (available in the HIPPIE2 software repository) using awk, bedtools v2.25.0 (Quinlan and Hall, 2010), and Python v2.7.9.

### Functional and genomic annotation data

We downloaded the cell type-specific ChIP-seq peak data for histone modifications (H3K4me1, H3K4me2, H3K4me3, H3K27ac, and H3K36me3), DNase I hypersensitive sites, and transcription factors or DNA-binding proteins (RNA Polymerase II, p300, and CTCF) from the 2011 freeze of the UCSC Genome Browser (Kent et al., 2002) for ENCODE datasets and directly from the web portal (https://egg2.wustl.edu/roadmap/web_portal/index.html) for Roadmap datasets including combinatorial epigenomic states from ChromHMM that we used to identify enhancer states (**Supplementary Table 1**).

### Enrichment analysis of functional genomic overlaps

To estimate the extent of overlap between PIRs and regulatory and epigenetic marks genome-wide, we calculated the sum of overlapped nucleotides between PIRs and each signal track (regulatory/epigenetic mark) genome-wide as the observed value. We sampled (1000 times) random genomic regions from the genome with length distribution matched with the length of PIRs. We calculated the average of 1,000 sums of overlaps between the sampled regions and each of the signal tracks. We then reported the percentage differences between the observed value and the averaged value from the background as the enrichment of the PIRs for each of the signal tracks. All region intersections were performed with bedtools v2.25.0 (Quinlan and Hall, 2010).

### Regulatory and genetic annotation of the interacting PIRs

HIPPIE2 annotates PIRs as enhancers, promoters, exons, introns, or intergenic elements. To do this, we used the cell-type-matched enhancer annotations described above and gene models downloaded from RefSeq (Pruitt et al., 2005). We annotate as enhancers the promoter-interacting PIRs that overlapped the enhancer (E) or weak enhancer (WE) annotation from the genome segmentation track (ChromHMM). We also annotated all promoter-interacting PIRs as an enhancer if they overlapped an open chromatin region with H3K4me1 or H3K27ac ChIP-seq peak, while not overlapping H3K4me3 and H3K27me3 peaks. The rest of the PIRs were annotated as promoters, exons, introns, and intergenic elements using RefSeq gene models (hg19 assembly). The promoters were defined as 500 bp-long regions upstream of the RefSeq TSS of protein-coding genes. We then annotated PIRs as promoters, exonic, intronic, or intergenic elements (in this prioritized order) based on their overlap with RefSeq gene models. To calculate the background expectations of interactions between annotations *a* and *b*, we used the product of the proportion of individual PIRs in annotation *a* and the proportion of those in annotation *b*.

### Transcription factor binding analysis of PIR–PIR interactions

To identify PIRs with evidence of transcription factor binding, we used Factorbook data (Wang et al., 2013) that integrates ChIP-seq experimental data from ENCODE with computationally-predicted TFBS to comprehensively survey protein–DNA binding genome-wide. The Factorbook data were obtained from UCSC hg19 database (factorbookMotifPos table, release 4). The Factorbook data contains 161 factors and the motifs were discovered from 91 cell types. We focused on 133 known DNA-binding transcription factors. We filtered out the TFs with less than 10 binding sites with PIRs genome-wide. For each PIR, we reported all TFs that have at least one binding site within that PIR. We reported enrichment for each of the surveyed binding motifs in PIR–PIR interactions. To do this, we categorized PIR–PIR interactions according to the classes of interacting PIR elements (enhancers, promoters, exons, introns, or intergenic elements). We estimated binding-motif enrichment as observed/expected frequency odds ratio. We computed the expected probability as the probability of the first class of PIR (C_i_) having one motif (M_k_) times the probability of the second class of PIR (C_j_) having another motif (M_l_) as follows:

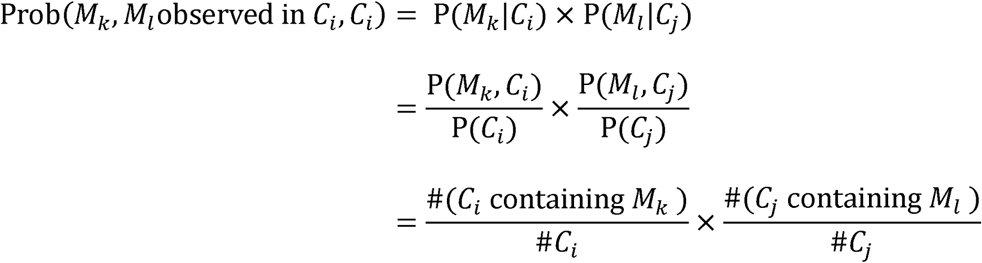

We performed a binomial distribution test to report the significance of observed binding motifs in each type of PIR–PIR interaction. To compare against the BioGRID database (Chatr-Aryamontri et al., 2015; Tyers et al., 2006), we downloaded the list of TF–TF interactions across cell lines and searched for transcription factor matches by name and by alias.

## Availability

HIPPIE2 software is freely available at https://bitbucket.org/wanglab-upenn/HIPPIE2. The corresponding interaction data tracks are available on the UCSC genome browser: https://genome.ucsc.edu/s/alexamlie/HIPPIE2%20vs%20Rao%20all%20cell%20lines%20darker%20interaction%20lines

## References

Ay, F., Bailey, T.L., and Noble, W.S. (2014). Statistical confidence estimation for Hi-C data reveals regulatory chromatin contacts. Genome Res. 24, 999–1011.

Beagan, J.A., Duong, M.T., Titus, K.R., Zhou, L., Cao, Z., Ma, J., Lachanski, C. V., Gillis, D.R., and Phillips-Cremins, J.E. (2017). YY1 and CTCF orchestrate a 3D chromatin looping switch during early neural lineage commitment. Genome Res. 27, 1139–1152.

Bernstein, B.E., Mikkelsen, T.S., Xie, X., Kamal, M., Huebert, D.J., Cuff, J., Fry, B., Meissner, A., Wernig, M., Plath, K., et al. (2006). A Bivalent Chromatin Structure Marks Key Developmental Genes in Embryonic Stem Cells. Cell 125, 315–326.

Bernstein, B.E., Birney, E., Dunham, I., Green, E.D., Gunter, C., and Snyder, M. (2012). An integrated encyclopedia of DNA elements in the human genome. Nature 489, 57–74.

Calo, E., and Wysocka, J. (2013). Modification of enhancer chromatin: what, how, and why? Mol. Cell 49, 825–837.

Chatr-Aryamontri, A., Breitkreutz, B.-J., Oughtred, R., Boucher, L., Heinicke, S., Chen, D., Stark, C., Breitkreutz, A., Kolas, N., O’Donnell, L., et al. (2015). The BioGRID interaction database: 2015 update. Nucleic Acids Res. 43, D470–8.

Consortium, R.E., Kundaje, A., Meuleman, W., Ernst, J., Bilenky, M., Yen, A., Heravi-Moussavi, A., Kheradpour, P., Zhang, Z., Wang, J., et al. (2015). Integrative analysis of 111 reference human epigenomes. Nature 518, 317–330.

Dang, C. V. (2012). MYC on the path to cancer. Cell 149, 22–35.

Dobin, A., Davis, C.A., Schlesinger, F., Drenkow, J., Zaleski, C., Jha, S., Batut, P., Chaisson, M., and Gingeras, T.R. (2013). STAR: ultrafast universal RNA-seq aligner. Bioinformatics 29, 15–21.

Durand, N.C., Shamim, M.S., Machol, I., Rao, S.S.P., Huntley, M.H., Lander, E.S., and Aiden, E.L. (2016). Juicer Provides a One-Click System for Analyzing Loop-Resolution Hi-C Experiments. Cell Syst. 3, 95–98.

Ernst, J., and Kellis, M. (2012). ChromHMM: automating chromatin-state discovery and characterization. Nat. Methods 9, 215–216.

Forcato, M., Nicoletti, C., Pal, K., Livi, C.M., Ferrari, F., and Bicciato, S. (2017). Comparison of computational methods for Hi-C data analysis. Nat. Methods 14, 679–685.

Heinz, S., Benner, C., Spann, N., Bertolino, E., Lin, Y.C., Laslo, P., Cheng, J.X., Murre, C., Singh, H., and Glass, C.K. (2010). Simple Combinations of Lineage-Determining Transcription Factors Prime cis-Regulatory Elements Required for Macrophage and B Cell Identities. Mol. Cell 38, 576–589.

Heinz, S., Romanoski, C.E., Benner, C., and Glass, C.K. (2015). The selection and function of cell type-specific enhancers. Nat. Rev. Mol. Cell Biol. 16, 144–154.

Hughes, J.R., Roberts, N., Mcgowan, S., Hay, D., Giannoulatou, E., Lynch, M., De Gobbi, M., Taylor, S., Gibbons, R., and Higgs, D.R. (2014). Analysis of hundreds of cisregulatory landscapes at high resolution in a single, high-throughput experiment. Nat. Genet. 46, 205–212.

Hwang, Y.-C., Zheng, Q., Gregory, B.D., and Wang, L.-S. (2013). High-throughput identification of long-range regulatory elements and their target promoters in the human genome. Nucleic Acids Res. 41, 4835–4846.

Hwang, Y.-C., Lin, C.-F., Valladares, O., Malamon, J., Kuksa, P., Zheng, Q., Gregory, B.D., and Wang, L.-S. (2014). HIPPIE: A high-throughput identification pipeline for promoter interacting enhancer elements. Bioinformatics 1–3.

Imakaev, M., Fudenberg, G., McCord, R.P., Naumova, N., Goloborodko, A., Lajoie, B.R., Dekker, J., and Mirny, L.A. (2012). Iterative correction of Hi-C data reveals hallmarks of chromosome organization. Nat. Methods 9, 999–1003.

Jin, F., Li, Y., Dixon, J.R., Selvaraj, S., Ye, Z., Lee, A.Y., Yen, C.-A., Schmitt, A.D., Espinoza, C.A., and Ren, B. (2013). A high-resolution map of the three-dimensional chromatin interactome in human cells. Nature 503, 290–294.

Kaplan, N., and Dekker, J. (2013). High-throughput genome scaffolding from in vivo DNA interaction frequency. Nat. Biotechnol. 31, 1143–1147.

Kent, W.J., Sugnet, C.W., Furey, T.S., Roskin, K.M., Pringle, T.H., Zahler, A.M., and Haussler, a. D. (2002). The Human Genome Browser at UCSC. Genome Res. 12, 996–1006.

Knight, P.A., and Ruiz, D. (2012). A fast algorithm for matrix balancing. IMA J. Numer. Anal. 33, 1029–1047.

Lajoie, B.R., Dekker, J., and Kaplan, N. (2015). The Hitchhiker’s guide to Hi-C analysis: practical guidelines. Methods 72, 65–75.

Lieberman-Aiden, E., van Berkum, N.L., Williams, L., Imakaev, M., Ragoczy, T., Telling, A., Amit, I., Lajoie, B.R., Sabo, P.J., Dorschner, M.O., et al. (2009). Comprehensive mapping of long-range interactions reveals folding principles of the human genome. Science (80-.). 326, 289–293.

Lun, A.T.L., and Smyth, G.K. (2015). diffHic: A Bioconductor package to detect differential genomic interactions in Hi-C data. BMC Bioinformatics 16, 1–11.

Ma, W., Ay, F., Lee, C., Gulsoy, G., Deng, X., Cook, S., Hesson, J., Cavanaugh, C., Ware, C.B., Krumm, A., et al. (2015). Fine-scale chromatin interaction maps reveal the cisregulatory landscape of human lincRNA genes. Nat. Methods 12, 71–78.

Mastrangelo, I.A., Courey, A.J., Wall, J.S., Jackson, S.P., and Hough, P. V (1991). DNA looping and Sp1 multimer links: a mechanism for transcriptional synergism and enhancement. Proc. Natl. Acad. Sci. 88, 5670–5674.

Mifsud, B., Tavares-Cadete, F., Young, A.N., Sugar, R., Schoenfelder, S., Ferreira, L., Wingett, S.W., Andrews, S., Grey, W., Ewels, P. a, et al. (2015). Mapping long-range promoter contacts in human cells with high-resolution capture Hi-C. Nat. Genet. 47, 598–606.

Norton, H.K., Emerson, D.J., Huang, H., Kim, J., Titus, K.R., Gu, S., Bassett, D.S., and Phillips-Cremins, J.E. (2018). Detecting hierarchical genome folding with network modularity. Nat. Methods 15, 119–122.

Ong, C.T., and Corces, V.G. (2014). CTCF: An architectural protein bridging genome topology and function. Nat. Rev. Genet. 15, 234–246.

Phillips, J.E., and Corces, V.G. (2009). CTCF: master weaver of the genome. Cell 137, 1194–1211.

Pruitt, K.D., Tatusova, T., and Maglott, D.R. (2005). NCBI Reference Sequence (RefSeq): a curated non-redundant sequence database of genomes, transcripts and proteins. Nucleic Acids Res. 33, D501–4.

Pruitt, K.D., Brown, G.R., Hiatt, S.M., Thibaud-Nissen, F., Astashyn, A., Ermolaeva, O., Farrell, C.M., Hart, J., Landrum, M.J., McGarvey, K.M., et al. (2014). RefSeq: An update on mammalian reference sequences. Nucleic Acids Res. 42, 756–763.

Quinlan, A.R., and Hall, I.M. (2010). BEDTools: a flexible suite of utilities for comparing genomic features. Bioinformatics 26, 841–842.

Rao, S.S.P., Huntley, M.H., Durand, N.C., Stamenova, E.K., Bochkov, I.D., Robinson, J.T., Sanborn, A.L., Machol, I., Omer, A.D., Lander, E.S., et al. (2014). A 3D Map of the Human Genome at Kilobase Resolution Reveals Principles of Chromatin Looping. Cell 159, 1665–1680.

Schreiber, J., Libbrecht, M., Bilmes, J., and Noble, W. (2018). Nucleotide sequence and DNaseI sensitivity are predictive of 3D chromatin architecture. BioRxiv 103614.

Shaulian, E., and Karin, M. (2002). AP-1 as a regulator of cell life and death. Nat. Cell Biol. 4, E131–E136.

Shlyueva, D., Stampfel, G., and Stark, A. (2014). Transcriptional enhancers: from properties to genome-wide predictions. Nat. Rev. Genet. 15, 272–286.

Thurman, R.E., Rynes, E., Humbert, R., Vierstra, J., Maurano, M.T., Haugen, E., Sheffield, N.C., Stergachis, A.B., Wang, H., Vernot, B., et al. (2012). The accessible chromatin landscape of the human genome. Nature 489, 75–82.

Tyers, M., Breitkreutz, A., Stark, C., Reguly, T., Boucher, L., and Breitkreutz, B.-J. (2006). BioGRID: a general repository for interaction datasets. Nucleic Acids Res. 34, D535–539.

Wang, J., Zhuang, J., Iyer, S., Lin, X.-Y., Greven, M.C., Kim, B.-H., Moore, J., Pierce, B.G., Dong, X., Virgil, D., et al. (2013). Factorbook.org: a Wiki-based database for transcription factor-binding data generated by the ENCODE consortium. Nucleic Acids Res. 41, D171–6.

Weintraub, A.S., Li, C.H., Zamudio, A. V., Sigova, A.A., Hannett, N.M., Day, D.S., Abraham, B.J., Cohen, M.A., Nabet, B., Buckley, D.L., et al. (2017). YY1 Is a Structural Regulator of Enhancer-Promoter Loops. Cell 171, 1573–1588.e28.

Whalen, S., Truty, R.M., and Pollard, K.S. (2016). Enhancer–promoter interactions are encoded by complex genomic signatures on looping chromatin. Nat. Genet. 48, 488–496.

Yaffe, E., and Tanay, A. (2011). Probabilistic modeling of Hi-C contact maps eliminates systematic biases to characterize global chromosomal architecture. Nat. Genet. 43, 1059–1065.

Yang, D., Jang, I., Choi, J., Kim, M.-S., Lee, A.J., Kim, H., Eom, J., Kim, D., Jung, I., and Lee, B. (2018). 3DIV: A 3D-genome Interaction Viewer and database. Nucleic Acids Res. 46, D52–D57.

Zhu, Y., Sun, L., Chen, Z., Whitaker, J.W., Wang, T., and Wang, W. (2013). Predicting enhancer transcription and activity from chromatin modifications. Nucleic Acids Res. 41, 10032–10043.

