## Supplementary Figures and Tables for "HIPPIE2: a method for fine-scale identification of physically interacting chromatin regions"

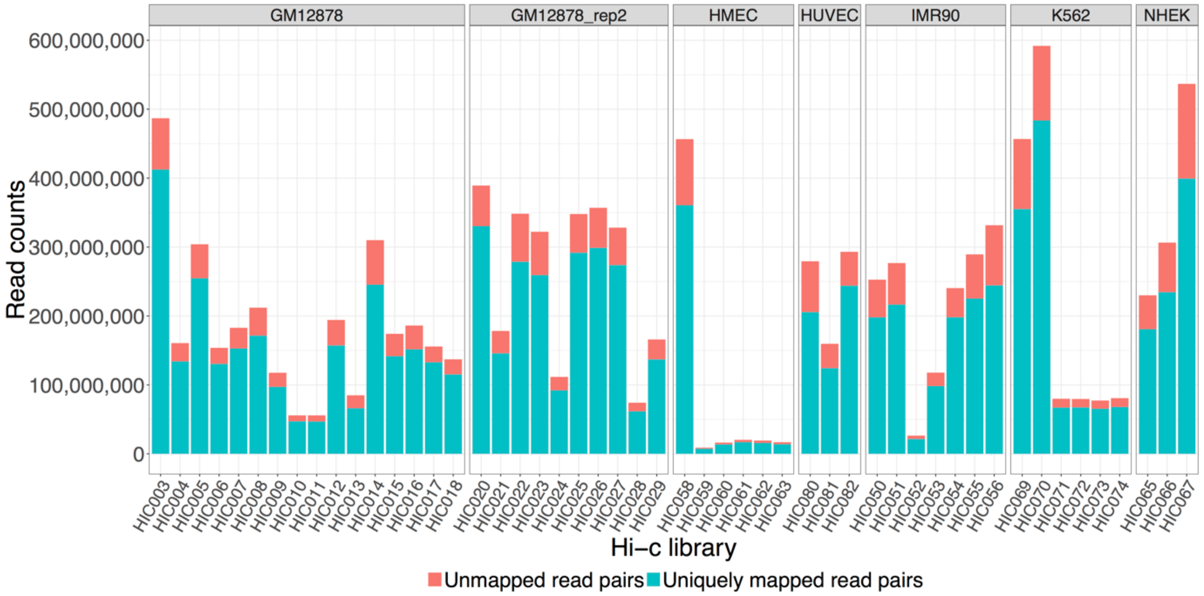

**Supplementary Figure 1.** Detailed HIPPIE2 mapping statistics for individual Hi-C libraries.

Uniquely mapped read pairs are used in HIPPIE2 analysis.

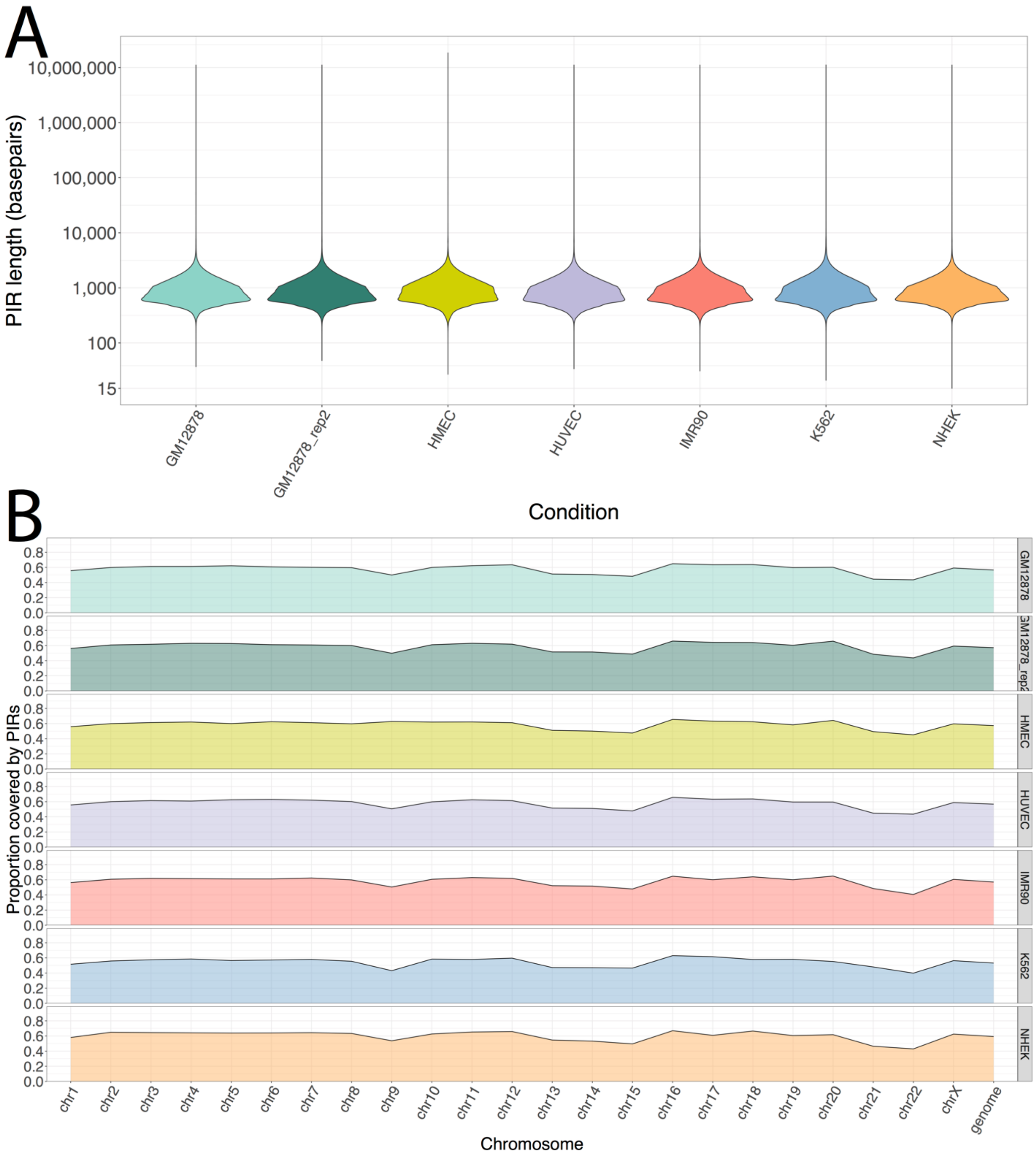

**Supplementary Figure 2. PIR length distributions.** A) Violin plots of PIR lengths across cell lines. B) PIR coverage patterns across chromosomes. 'Genome" refers to total genomic coverage.

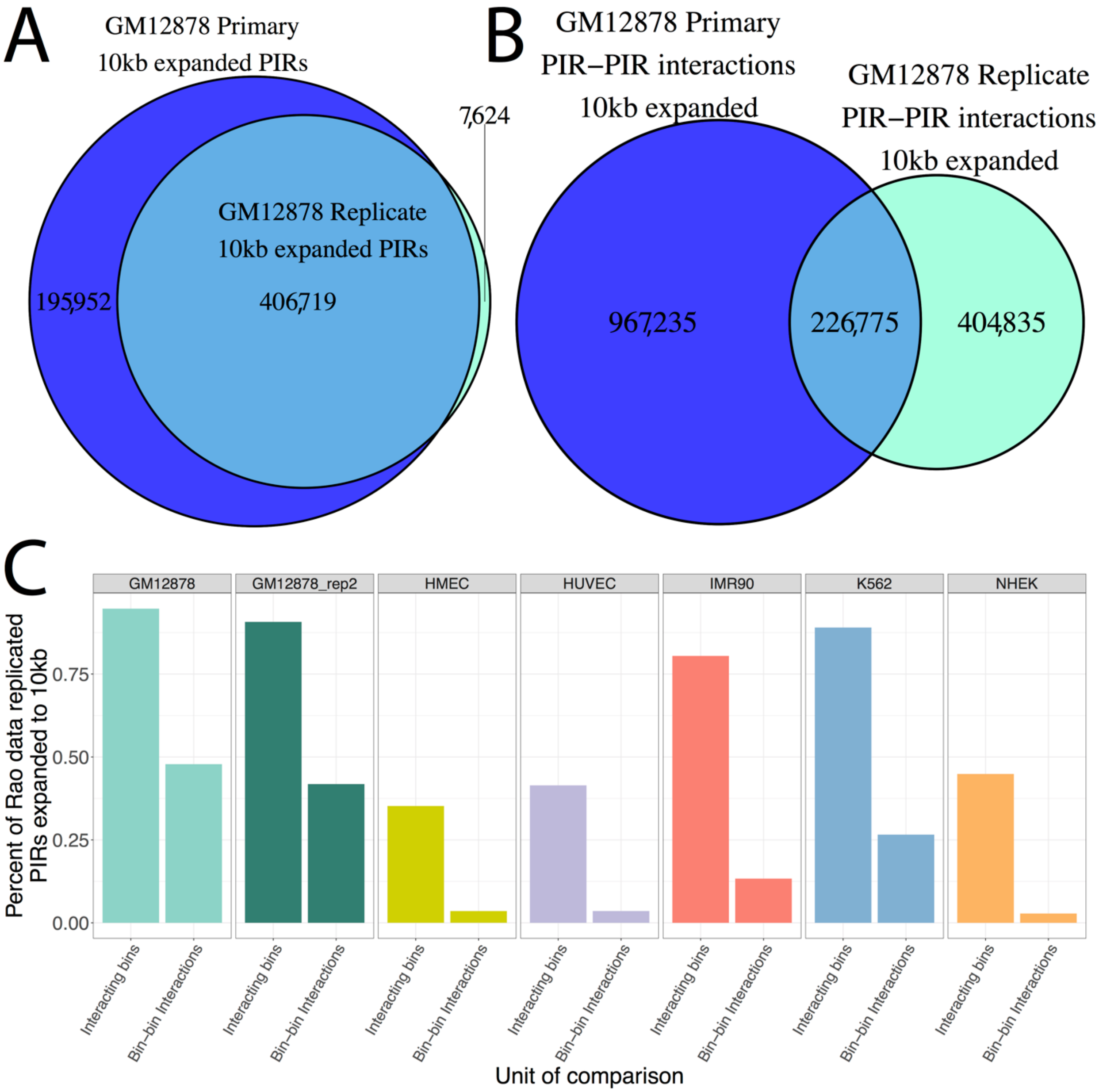

**Supplementary Figure 3. 10kb expanded PIR comparisons.** A) Replication of PIRs involved in significant interactions, expanded to at least 10 kilobases between the primary and secondary GM12878 libraries. B) Replication of PIR–PIR interactions which each side expanded to at least 10 kilobases between the primary and secondary GM12878 libraries. C) Replication rates of Rao data by HIPPIE2 PIRs expanded to 10kb.

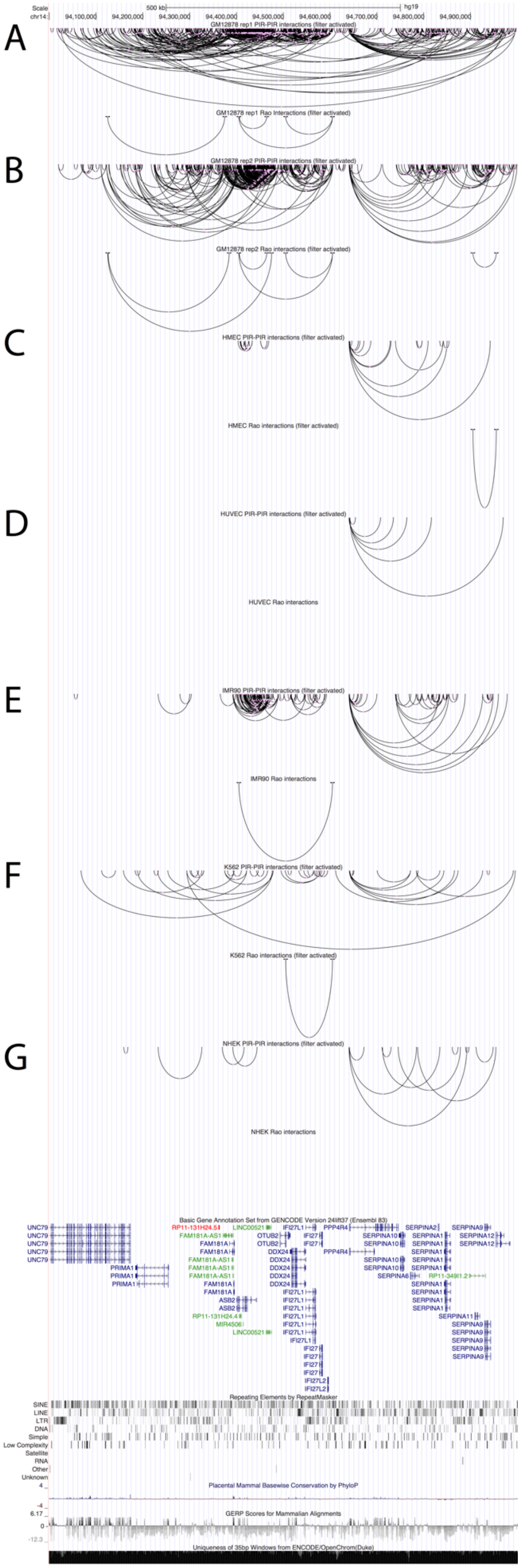
A
B
C
D
E
F
G
GM12878 rep1 PIR-PIR interactions (filter activated)
GM12878 rep1 Rao interactions (filter activated)
GM12878 rep2 PIR-PIR interactions (filter activated)
GM12878 rep2 Rao interactions (filter activated)
HMEC PIR-PIR interactions (filter activated)
HMEC Rao interactions (filter activated)
HUVEC PIR-PIR interactions (filter activated)
HUVEC Rao interactions
IMR90 PIR-PIR interactions (filter activated)
IMR90 Rao interactions
K562 PIR-PIR interactions (filter activated)
K562 Rao interactions (filter activated)
NHEK PIR-PIR interactions (filter activated)
NHEK Rao interactions
Basic Gene Annotation Set from GENCODE Version 24lift37 (Ensembl 83)
Repeating Elements by RepeatMasker
Placental Mammal Basewise Conservation by PhyloP
GERP Scores for Mammalian Alignments
Uniqueness of 35bp Windows from ENCODE/OpenChrom(Duke)

**Supplementary Figure 4. Genome browser shot of HIPPIE2 vs Rao data.** A) GM12878 primary HIPPIE2 vs Rao signals. B) GM12878 replicate library HIPPIE2 vs Rao signals. C) HMEC HIPPIE2 vs Rao signals. D) HUVEC HIPPIE2 vs Rao signals. E) IMR90 HIPPIE2 vs Rao signals. F) K562 HIPPIE2 vs Rao signals. G) NHEK HIPPIE2 vs Rao signals.

 

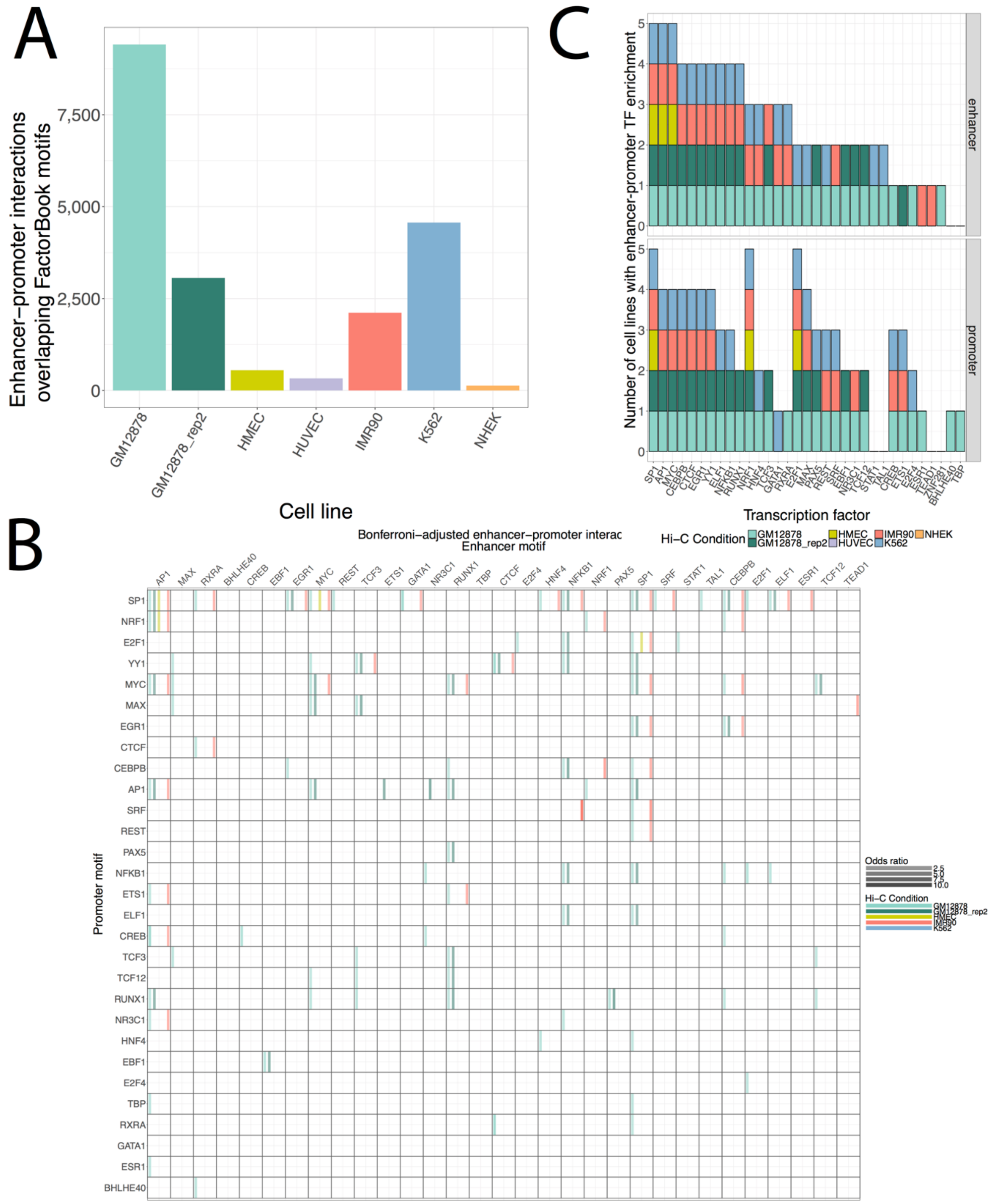

**Supplementary Figure 5. TFBS analysis of HIPPIE2 interactions.** A) Number of interactor PIRs overlapping FactorBook transcription factor binding sites. B) Heatmap of odds ratio of observed TF–TF interactions relative to background expectation calculated by genomic coverage of TFBSs. C) Transcription factor enrichmen patternst in enhancer and promoter elements across cell lines.

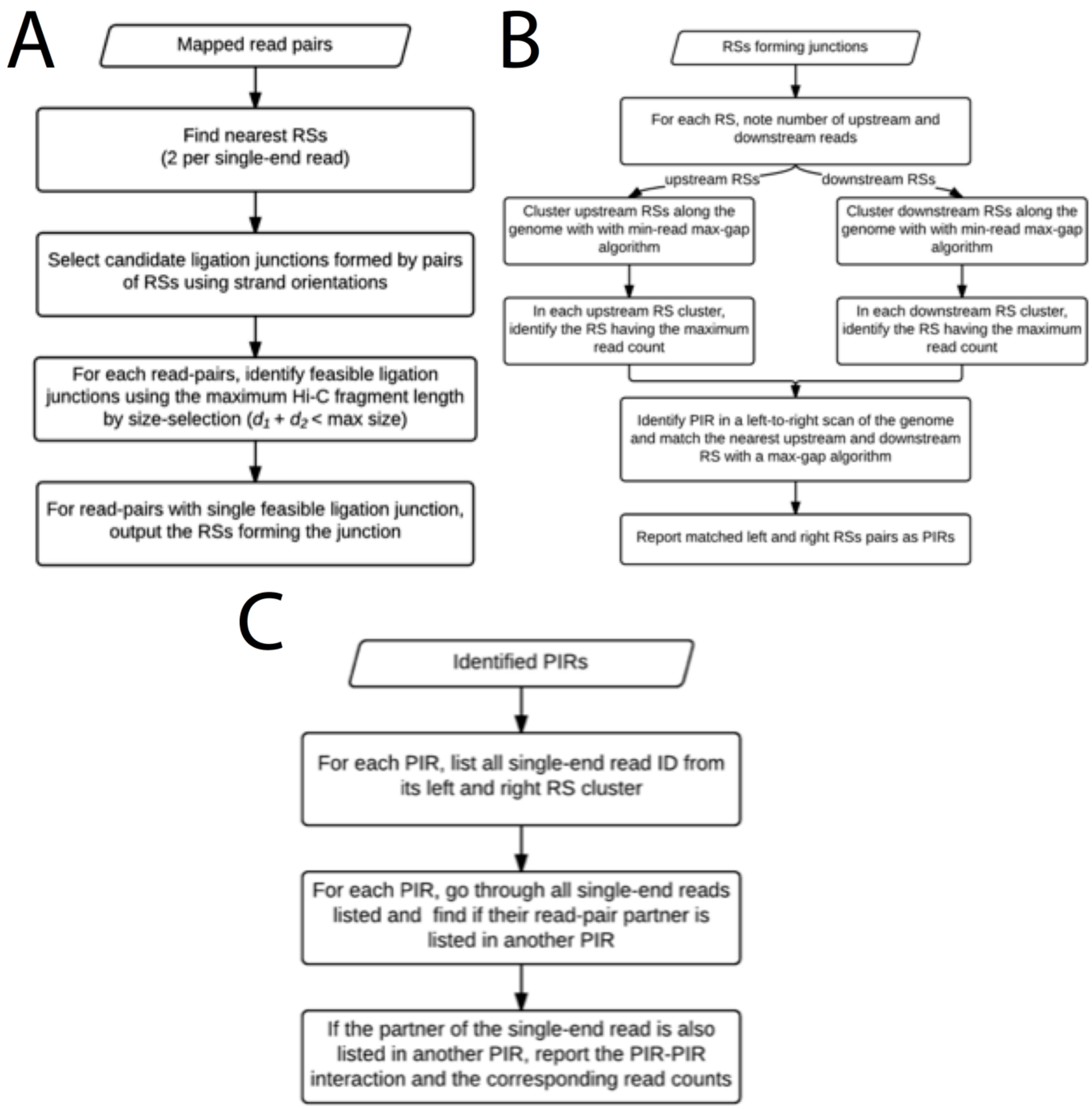

**Supplementary Figure 6. Flowchart of HIPPIE2 PIR identification and PIR-PIR interaction identification approach.** A) Identifying restriction sites (RSs) corresponding to cut/ligation events. B) Identifying PIR left and right boundaries using max-gap/min-height clustering of cut/ligation sites. C) Identifying PIR-PIR interactions.

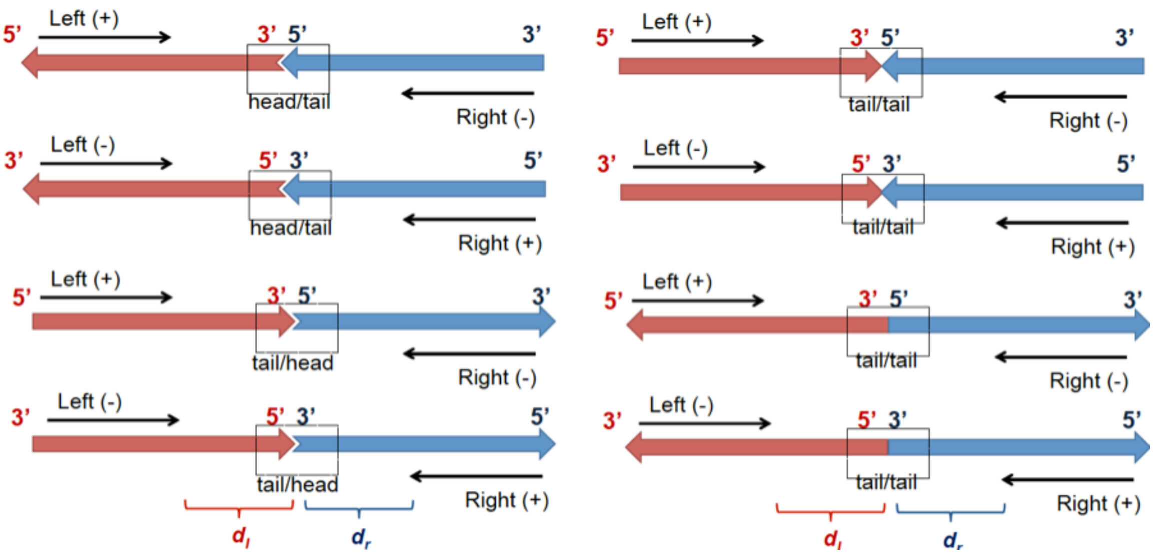

**Supplementary Figure 7. DNA-DNA ligation configurations.** (Left) Four possible ligation scenarios for the strand orientation of the read-pair as +/- or -/+ (different strand; case 1). (Right) Four possible ligation scenarios for the read-pair with strand orientation +/+ or -/- (same strand; case 2). Black arrows: sequencing read orientation; red and blue boxes with arrowheads: ligated DNA fragments and their coordinate orientations; dotted box: the ligation type for the orientation for the ends from both ligated DNA fragments.

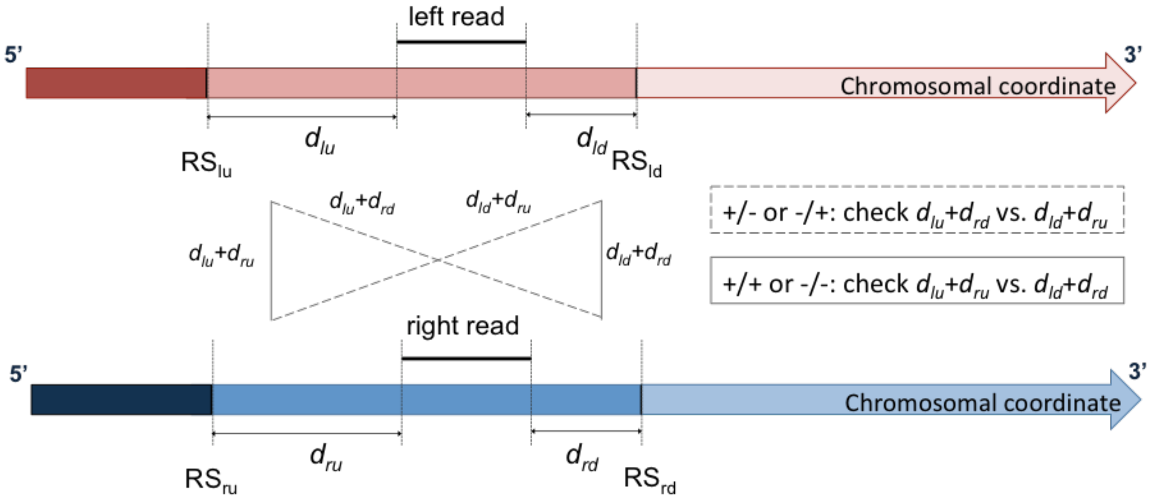

**Supplementary Figure 8. Strategy for determining the cut-and-ligation sites** for read-pair with different strand (case 1) or same strand (case 2). $RS_{lu}$: the nearest upstream restriction site from the left read; $RS_{ld}$: the nearest downstream restriction site from the left read; $RS_{ru}$: the nearest upstream restriction site from the right read; $RS_{rd}$: the nearest downstream restriction site from the right read; $d_{lu}$: the distance from the left read upstream end to $RS_{lu}$; $d_{ld}$: the distance from the left read downstream end to $RS_{ld}$; $d_{ru}$: the distance from the right read upstream end to $RS_{ru}$; $d_{rd}$: the distance from the right read downstream end to $RS_{rd}$.

### Supplementary Tables

#### Supplementary Table 1. Functional and genomics data integrated with HIPPIE2 (hippie2_suppl_table_1.xlsx)

#### Supplementary Table 2. Strand percentage of Hi-C read pairs

| All read pairs | | | |
|---|---|---|---|
| read1 | read2 | count | % |
| + | + | 351,356,711 | 20.4 |
| + | - | 507,746,008 | 29.5 |
| - | + | 505,952,205 | 29.4 |
| - | - | 355,039,391 | 20.6 |
| Read pair distance >= 5000 bp | | | |
| read1 | read2 | count | % |
| + | + | 329,957,629 | 24.9 |
| + | - | 331,958,821 | 25.0 |
| - | + | 330,499,655 | 24.9 |
| - | - | 333,438,805 | 25.1 |

**Supplementary Table 3: Enrichment patterns of HIPPIE2-identified transcription factors in enhancer–promoter interactions across cell lines**

| Transcription Factor | Enhancer cell line enrichments | Promoter cell line enrichments |
|---|---|---|
| SP1 | GM12878; GM12878_rep2; HMEC; IMR90; K562 | GM12878; GM12878_rep2; HMEC; IMR90; K562 |
| AP1 | GM12878; GM12878_rep2; HMEC; IMR90; K562 | GM12878; GM12878_rep2; IMR90; K562 |
| MYC | GM12878; GM12878_rep2; HMEC; IMR90; K562 | GM12878; GM12878_rep2; IMR90; K562 |
| CEBPB | GM12878; GM12878_rep2; IMR90; K562 | GM12878; GM12878_rep2; IMR90; K562 |
| CTCF | GM12878; GM12878_rep2; IMR90; K562 | GM12878; GM12878_rep2; IMR90; K562 |
| EGR1 | GM12878; GM12878_rep2; IMR90; K562 | GM12878; GM12878_rep2; IMR90; K562 |
| YY1 | GM12878; GM12878_rep2; IMR90; K562 | GM12878; GM12878_rep2; IMR90; K562 |
| ELF1 | GM12878; GM12878_rep2; IMR90; K562 | GM12878; GM12878_rep2; K562 |

| | | |
|---|---|---|
| NFKB1 | GM12878; GM12878_rep2; IMR90; K562 | GM12878; GM12878_rep2; K562 |
| RUNX1 | GM12878; GM12878_rep2; IMR90; K562 | GM12878; GM12878_rep2 |
| NRF1 | GM12878; IMR90; K562 | GM12878; GM12878_rep2; HMEC; IMR90; K562 |
| HNF4 | GM12878; IMR90; K562 | GM12878; K562 |
| TCF3 | GM12878; GM12878_rep2; IMR90 | GM12878; GM12878_rep2 |
| GATA1 | GM12878; IMR90; K562 | K562 |
| RXRA | GM12878; IMR90; K562 | GM12878 |
| STAT1 | GM12878; K562 | None |
| TAL1 | GM12878; K562 | None |
| E2F1 | GM12878; K562 | GM12878; GM12878_rep2; HMEC; IMR90; K562 |
| MAX | GM12878; K562 | GM12878; GM12878_rep2; IMR90; K562 |
| PAX5 | GM12878; GM12878_rep2 | GM12878; GM12878_rep2; K562 |
| REST | GM12878; K562 | GM12878; IMR90; K562 |
| SRF | GM12878; IMR90 | GM12878; IMR90; K562 |

| | | |
|---|---|---|
| EBF1 | GM12878; GM12878_rep2 | GM12878; GM12878_rep2 |
| NR3C1 | GM12878; GM12878_rep2 | GM12878; IMR90 |
| TCF12 | GM12878; GM12878_rep2 | GM12878; GM12878_rep2 |
| TEAD1 | IMR90 | None |
| ZNF281 | GM12878 | None |
| CREB | GM12878 | GM12878; IMR90; K562 |
| ETS1 | GM12878_rep2 | GM12878; IMR90; K562 |
| E2F4 | GM12878 | GM12878; K562 |
| ESR1 | IMR90 | GM12878 |
| BHLHE40 | None | GM12878 |
| TBP | None | GM12878 |
